# Development and evaluation of a rapid on-site water pathogen detection system for water quality monitoring

**DOI:** 10.1101/731778

**Authors:** Joel Y. Chia, Yu Pei Tay, Daniel Poh, Boon Hunt Tay, Eileen Koh, Joshua Teo, Hoi Ming Yap, Merrill Lim, Li Ting Soh, Yuan Kun Lee, Chi-Lik Ken Lee

## Abstract

Timely response to outbreak of water-borne diseases caused by bacteria requires efficient monitoring and rapid detection strategies. Herein, we report a rapid DNA-based on-site detection system for specific detection of *Pseudomonas aeruginosa.* To evaluate the test performance of our method against spiked water samples, parallel tests based on real-time PCR and standard culture methods were concurrently performed. Test sensitivities of between 96.7% and 92.3% were obtained, based on the calculation obtained from qPCR and culture test, respectively, with a corresponding level of specificity of 92.9% and 83.3%. Time-to-result is around 45 min, with a detection limit of 1 CFU/100 mL. Here, a fully-deployable detection method where bacteria of-interest can be detected rapidly with high accuracy was described. This test method can be modified to detect other bacteria of-interest and can also be used in different applications. The test results can be obtained on-site and can therefore be particularly useful in public facilities health surveillance, where regulators can quickly determine if a site is safe or if other emergency response measures are required.

**IMPORTANCE:** *Pseudomonas aeruginosa* is a well-established water quality biomarker, known to be associated to humans’ health risks. To ascertain the presence of this pathogen, relevant stakeholders currently rely on standard quantitative ISO methods (APHA 9213E) which require 6 days from sampling to results. This window could potentially lead to waterborne outbreaks if the contaminated water features are not shut down for proper and urgent mitigation. This manuscript describes a method to detect this disease-causing microorganism in its viable forms under 1 hour, with a sensitivity of 1 CFU/100mL. Besides providing valuable information of the quality of water system, this direct monitoring of pathogens can reduce substantial time needed from sampling to reporting. This method can be established as a platform technology for other pathogenic microorganisms. On-going work to develop economic point-of-care prototypes could facilitate quick screening of targeted waterborne pathogens and results in better assessment of public health risk and quick in devising emergency response measures and other management strategies.

## INTRODUCTION

Clean and safe water are essential to public health, and the ability to rapidly monitor its quality and detect for the presence of water pathogens are indispensable. Currently, the standard culture method remains the most widely used method, despite the fact that the time required from sample to results can often take as long as from a few days to close to 2 weeks. Moreover, the results obtained often might not be as reliable as might be desired and that only a very small fraction (0.01 - 1 %) of viable bacteria can be detected (Köster et al. 2003; Watkins and Jian 1997). This is further complicated by the fact that certain microorganisms such as *Mycobacterium tuberculosis* and *Legionella pneumophila* are known to be either difficult to culture *in vitro* and/or are known to remain in a viable but nonculturable state (VBNC) (Casini et al. 2017; Keserue et al. 2012; Dunn, Starke, and Revell 2016). In addition, there are also a number of intrinsic problems and limitations associated with the pour plate, spread plate and membrane filtration techniques involved in the traditional cultivation method (Köster et al. 2003).

Recognising the limitations of the culture methods, the U.S. Environmental Protection Agency (EPA) has adopted qPCR as a recommended and alternative method for rapidly enumerating *Enterococcus* species in recreational surface water (‘Recreational water quality criteria. Technical Report EPA-820-F-12-058; Office of Water, U.S. Environmental Protection Agency: Washington, DC.’ 2012). Results can be obtained within 3 to 4 hours as opposed to 18 to 72 hours for the culture method for the same bacteria. During the process, the target of interest (i.e. specific DNA from the bacteria of interest) is exponentially amplified, allowing for the detection of the target, even at very low initial concentration. The challenge now, is to implement qPCR as an on-site surveillance method where public health officers can easily and quickly assess water quality and decision can be made, with the necessary course of action timely executed. This can help minimise unnecessary shut-down of facilities while not jeopardising public health. Although significant advances have been made in developing fully integrated and automated portable PCR devices, the implementation of these platforms are still let down by their cost effectiveness (Quan, Sauzade, and Brouzes 2018; Zhang and Jiang 2016; Huggett et al. 2008). These have led to the emergence of alternative technology such as the various isothermal DNA amplification approaches where the reaction is carried out at a constant (and often lower) temperature, which completely eliminates the need for thermal cycling (and the additional time needed for cooling and heating) and the requirement for costly temperature controller. To date, isothermal DNA amplification has shown to be very promising, especially in terms of their reliabilities, amplification efficiencies, robustness, speed, and cost effectiveness (Quan, Sauzade, and Brouzes 2018; Lau and Botella 2017; Kolm et al. 2017). However, to our knowledge, most if not all of the developments are still rudimentary in terms of their deployability, where additional lab-based equipment and extraction protocols are still needed (Kolm et al. 2017; Wong et al. 2018; Fuller et al. 2017; Jauset-Rubio et al. 2016; Lalle et al. 2018; Wang et al. 2017; Mamba et al. 2018; Organization 2016).

Here, we described the development of a fully deployable isothermal DNA amplification detection method for water microbial quality testing, utilising the recombinase polymerase isothermal technology (TwistDx Inc., UK). Test were performed and the results were referenced against the parallel test results obtained from the standard culture method and qPCR. We were able to achieve a level of sensitivity and specificity of 96.7% and 92.9% respectively (using qPCR as reference) and a corresponding test performance of 92.3% (sensitivity) and 83.3% (specificity), with the cell culture method as reference.

## MATERIALS AND METHODS

### Primer and probe design

The sequences for the qPCR primers (forward primer: 5’-TTCCCTCGCAGAGAAAACATC-3’; reverse primer: 5’-CCTGGTTGATCAGGTCGATCT-3’) targeting the *algD* GDP mannose dehydrogenase gene of *Pseudomonas aeruginosa (P. aeruginosa),* GenBank ref. Y00337.1, were obtained from da Silva Filho *et al.* (da Silva Filho et al. 1999). The sequences for the flanking primers for RPA (forward primer: 5’-TTCCCTCGCAGAGAAAACATCCTATCACCGCG-3’; reverse primer: 5’-Biotin-GGGCGACTTGCCCTGGTTGATCAGGTCGATCT-3’) were derived and modified from the original sequences (da Silva Filho et al. 1999). The internal probe sequence (5’-FAM-CGGCCACCTCATTAACGGCGCGACAAACAATC-dSpacer-AGGTGAATGCGATGC-3’-phosphorylation) were obtained from a suitable internal sequences within the region flanked by the flanking primers. The sequences for the qPCR primers for Enterococcus species 16S rRNA (forward primer: 5’-TGCATTAGCTAGTTGGTG-3’; reverse primer: 5’-TTAAGAAACCGCCTGCGC-3’) were obtained from published report(Ryu et al. 2012).

### Preparation of cell lysate

*P.aeruginosa* (ATCC 27853) was cultured overnight in nutrient broth at 37°C with shaking. A 10-fold serial dilution was performed on the subsequent day, and aliquots from each dilution were taken out and subjected to heat lysis at 95°C for 10 min. The cell lysates will be used for both the qPCR and RPA experiment. In order to quantify the amount of viable cells originally present in each of the various dilutions, aliquots from the original diluted and unheated samples were separately plated on nutrient agar and incubated overnight for numeration of the colonies on the following day.

### qPCR analysis

qPCR was performed using the iTaq Universal SYBR Green Supermix (Bio-Rad, Hercules, CA), in the presence of the forward and reverse primers ([500 nM]_final_, each) and 2 uL of *P.aeruginosa* lysate, in a reaction volume of 20 uL. The amplification was carried out for 40 cycles of 95°C for 3 sec and 60°C for 1 min after an initial heating at 95°C for 4 min using the Bio-Rad CFX-96 real-time PCR system. A melt curve analysis from 65°C from 95°C was also performed.

### Recombinase Polymerase amplification

RPA reaction were performed using the TwistAmp nfo kit (TwistDx Inc.) as according to the manufacturer’s protocol. The reaction product was diluted and analysed using the HybriDetect (Milenia Biotec, Giessen, Germany) lateral kit /dipstick as according to the manufacturer’s instruction. The detailed test procedure is currently not available^*^.

### Calculation of test performance

The number of true positive (a), false positive (b), false negative (c) and true negative (d) test results were tallied and the test performance was calculated based on the following formulas. Test sensitivity= a/(a+c); test specificity= d/(b+d); positive predictive value (PPV)= a/(a+b) and negative predictive value (NPV)= d/(c+d).

## RESULTS AND DISCUSSION

### Validation of DNA primers specificity

DNA-based amplification detection methods allow for the amplification and detection of the specific target, such as bacteria of interest. To achieve this, target-specific DNA primers and probe are needed. To validate the specificity of the DNA primers used in our test, we performed qPCR against varying concentrations of both the positive and the negative control targets. Figure 1A shows the amplification plot of the reactions, where the increase in fluorescence corresponds to the increase in amplification products. As expected, no amplification products were detected from the no-template control and from all the negative control reactions. All the reactions where the positive control targets are present showed positive signals. As a quality control test, the same set of samples were tested using the primers specific for the original negative control (Figure 1B). It is important to note that the DNA-based amplification method, besides its sensitivity, has an added advantage over antibodies-based method, in terms of its versatility and ease to detect for different targets and variants, through changing the primary sequence of the primers and probe.

**Figure 1.**
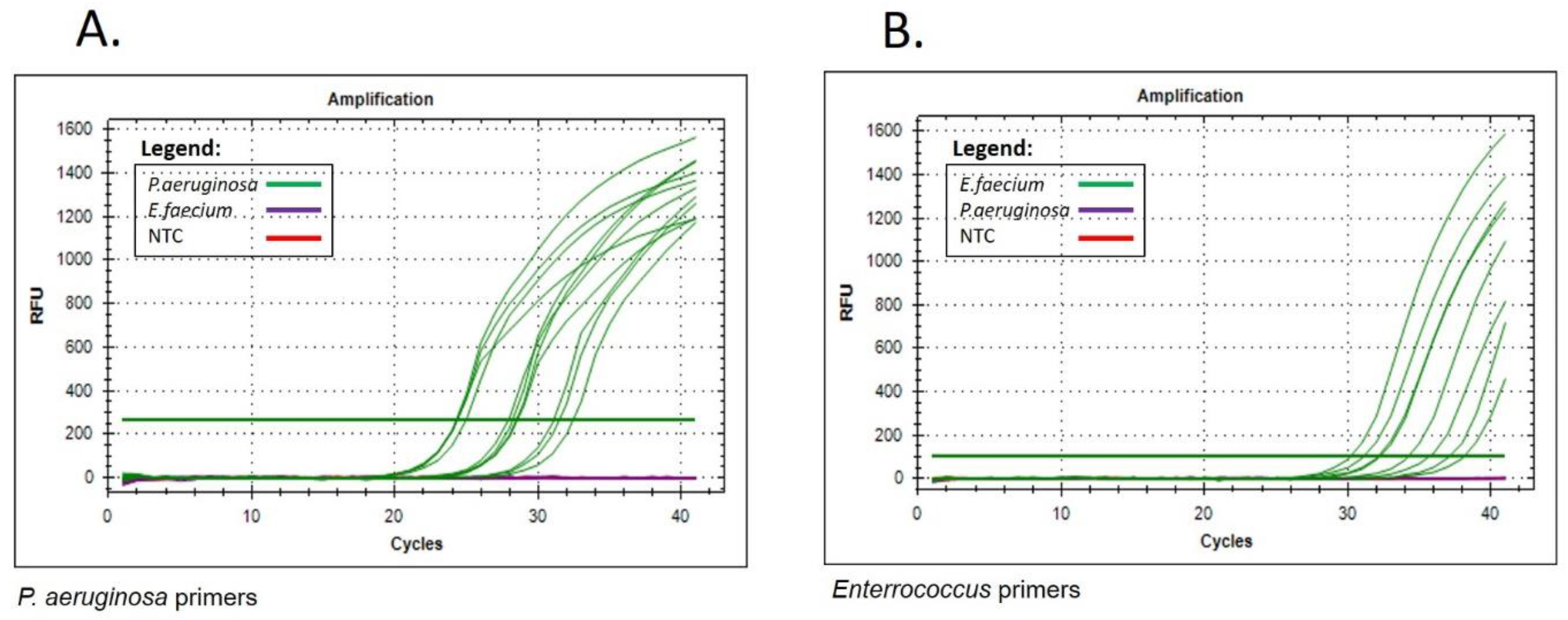
Validation of primers specificity A) The qPCR amplification plot showing the PCR reaction where the specificity of the *P.aeruginosa* primers are tested against *P.aeruginosa* (positive control), *E.faecium* (negative control) and no-template control. Decreasing amount of *P.aeruginosa* (2800 CFU, 280 CFU and 28 CFU) and *E.faecium* (5100 CFU, 510 CFU and 51 CFU) was used as the template in each reaction well. A minimum of triplicates were performed for each concentration of samples. B) The qPCR amplification plot showing the PCR reaction using the *Enterococcus* primers against the same set of samples.

### Amplification of target bacteria DNA via isothermal DNA amplification

To address the need for a rapid, sensitive and fully deployable water testing kit where presence of specific bacteria of interest can be reliably detected, isothermal recombinase polymerase DNA amplification (RPA) and lateral flow immunoassay were adopted in our test design (TwistDx Inc., UK.). Figure 2A is a schematic diagram depicting the principles and mechanism of detection. Briefly, DNA from the bacteria of interest is amplified via RPA and doubly-labelled using the relevant antigenic and detection tags. The amplification product is subsequently captured and detected using a customised lateral flow kit. The double-labelling approach ensures only the correct (i.e. positive) amplicons can be captured and detected by the lateral flow kit, reducing the likelihood of a false positive readout.

**Figure 2:**
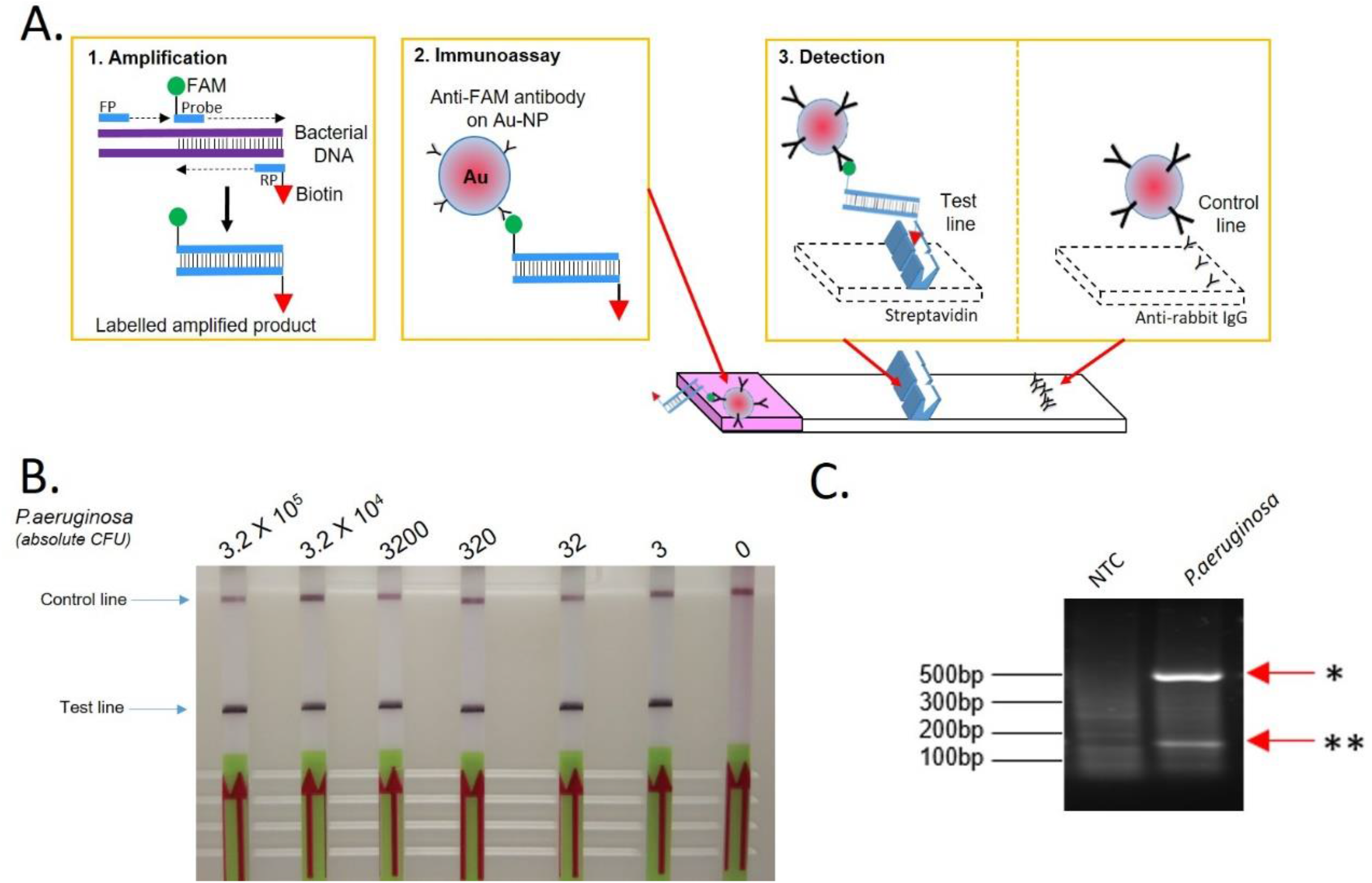
Amplification of target bacteria DNA via isothermal DNA amplification and detection via lateral flow immunoassay A) Schematic diagram depicting the mechanism of detection. Step 1: DNA from the bacteria of interest is subjected to isothermal recombinase polymerase DNA amplification using specific oligo-probe and oligo-primers, together will all the necessary enzymes, reactants and buffered solution at 37°C for 30-40 min. Step 2: The amplification product mix is applied to a customised lateral flow kit, where the amplicons can be captured by the anti-FAM conjugated gold-nano particles. Step 3: The complex, comprising of the amplicon and the anti-FAM gold-nano particle flows along the lateral kit and can then be captured by immobilised streptavidin or anti-biotin (at the “Test line” position), resulting in the aggregation and coloured visualisation of the complex. Excess or unreacted anti-FAM conjugated gold-nano particles will be captured and visualised at the “Control line” position. In the absence of the amplicon (i.e. a negative test), the lateral kit will only shows one band at the “Control line” position. In the presence of the amplicon (i.e. a positive test), the lateral kit will show two bands, one at the “Test line” and another at the “Control line” position. Note: the “Control line” serves as an internal control, to indicate that the lateral immunoassay has been completed and is successful, and needs to show up for every valid test. If it is absent, the test for this specific lateral kit is considered invalid. Legend: FP: forward primer; RP: reverse primer; FAM: Fluorescein probe/tag B) Different amount of *P.aeruginosa* is prepared and heat-lysed and specific amount of the lysates are subjected to isothermal recombinase polymerase amplification and the amplification products subjected to detection via the lateral test kit immunoassay as described above. Standard culture plate count for the *P.aeruginosa* preparation were performed concurrently, so as to ascertain the actual amount of *P.aeruginosa* in the test samples, as per during the test. Shown here is the test result and the actual amount of *P.aeruginosa* as per during the day of the test, in decreasing amount. All the samples that contain *P.aeruginosa* yielded positive test results (from the highest, to the lowest amount of *P.aeruginosa* in this batch of samples). In the absence of *P.aeruginosa* (i.e 0 CFU), there was no band observed at the “Test line”, correctly indicating the absence of *P.aeruginosa.* C) Agarose gel electrophoresis (AGE) showing the amplification products from RPA. The amplification products were purified prior to AGE using the Promega PCR Clean-up kit. Legend: NTC: No-template control; *: first amplification product; **: second amplification product.

Figure 2B shows the test results for the amplification of *P.aeruginosa* DNA by RPA. Presence of a band at the test line position indicate a positive test result. All the samples that contain *P.aeruginosa* returned positive test results while the control sample that did not contain *P.aeruginosa* also correctly shows the absence of *P.aeruginosa* as shown by the absence of a band at the test line position. All the samples used were plated and quantified using standard culture plate count (data not shown) on the same day of the test. Based on the amount used in the experiment, we were able to detect as low as 3 CFU^†^ of *P.aeruginosa.* To verify the test result, the amplification product (obtained from the sample that contains 3 CFU of *P.aeruginosa)* was analysed using agarose gel electrophoresis. Figure 2C shows the presence of the amplification products of the expected and correct size. To further investigate the detection limit of the assay, additional RPA test using concentration down to 1 CFU *P.aeruginosa* (Figure 3A), together with qPCR as a parallel test were also performed (Figure 3B). The results show that we were able to detect down to 1 CFU.

**Figure 3:**
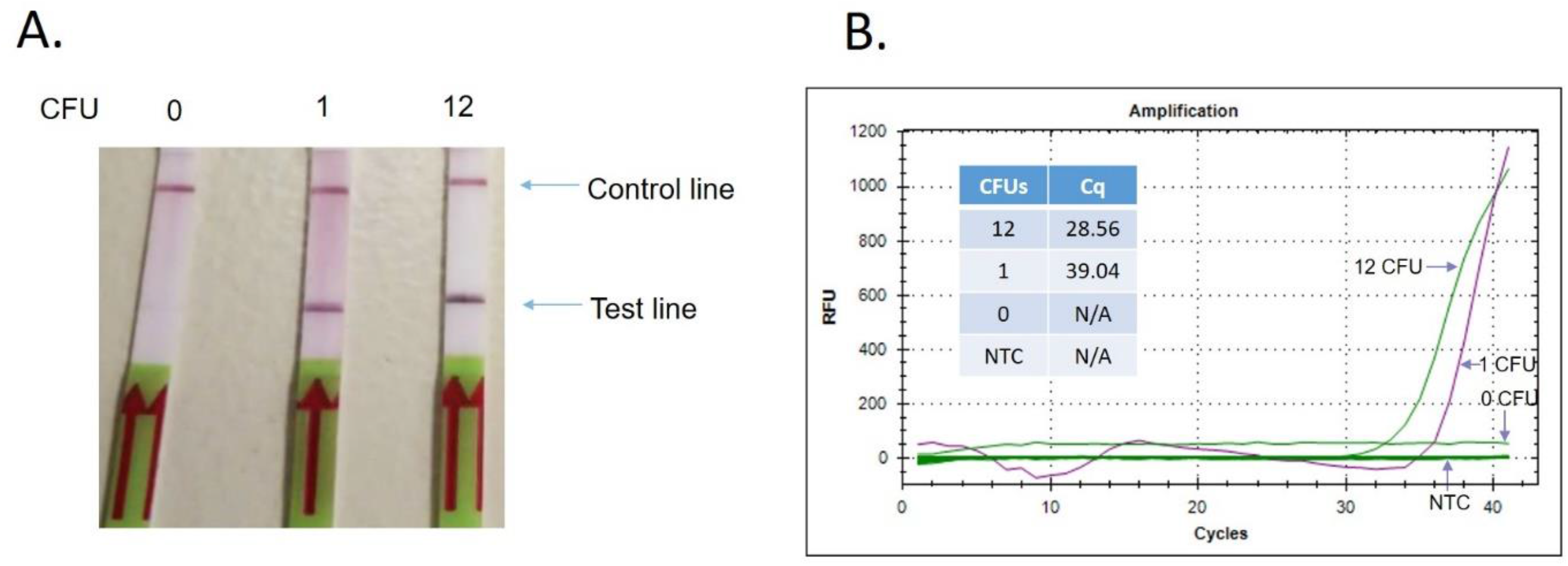
Detection using lower amount of samples A) Different amount of *P.aeruginosa* is prepared and heat-lysed and specific amount of the lysates are subjected to isothermal recombinase polymerase amplification and the amplification products subjected to detection via the lateral test kit immunoassay as aforementioned. B) The qPCR amplification plot showing the PCR reaction using the *P.aeruginosa* primers against the same set of samples. The CFU equivalent amount of samples and the Cq (Cycle threshold) are shown in the inset table.

### Detection using spiked samples

To further evaluate the RPA-lateral flow detection system as a feasible water test system, a complete water testing work flow was designed and tested against water spiked with *P.aeruginosa.* Figure 4A outlines the workflow, which includes: i) a filtering/concentration step; ii) a extraction/recovery step; and iii) a amplification/detection step. Based on this workflow, different amount of *P.aeruginosa* was separately spiked into 250-mL of sterile ultra-pure water and tested as described. Figure 4B is a representative data taken from a series of test performed (n=90). Table 1 shows the overall test performance. The calculation was done based on qPCR and standard culture plate count each as the separate standard reference, and the number of positive, negative, false positive and false negative results tallied. Here, two set of data were obtained. In summary, the test obtained a level of sensitivity of 96.7% and 92.3%, based on the calculation obtained from qPCR and culture test, respectively. Correspondingly, a level of specificity of 92.9% and 83.3% were also obtained. The probability to correctly screen out positive or negative samples (i.e. PPV and NPV) also returned highly favourable value of between 83.3% to 96.7%.

**Figure 4:**
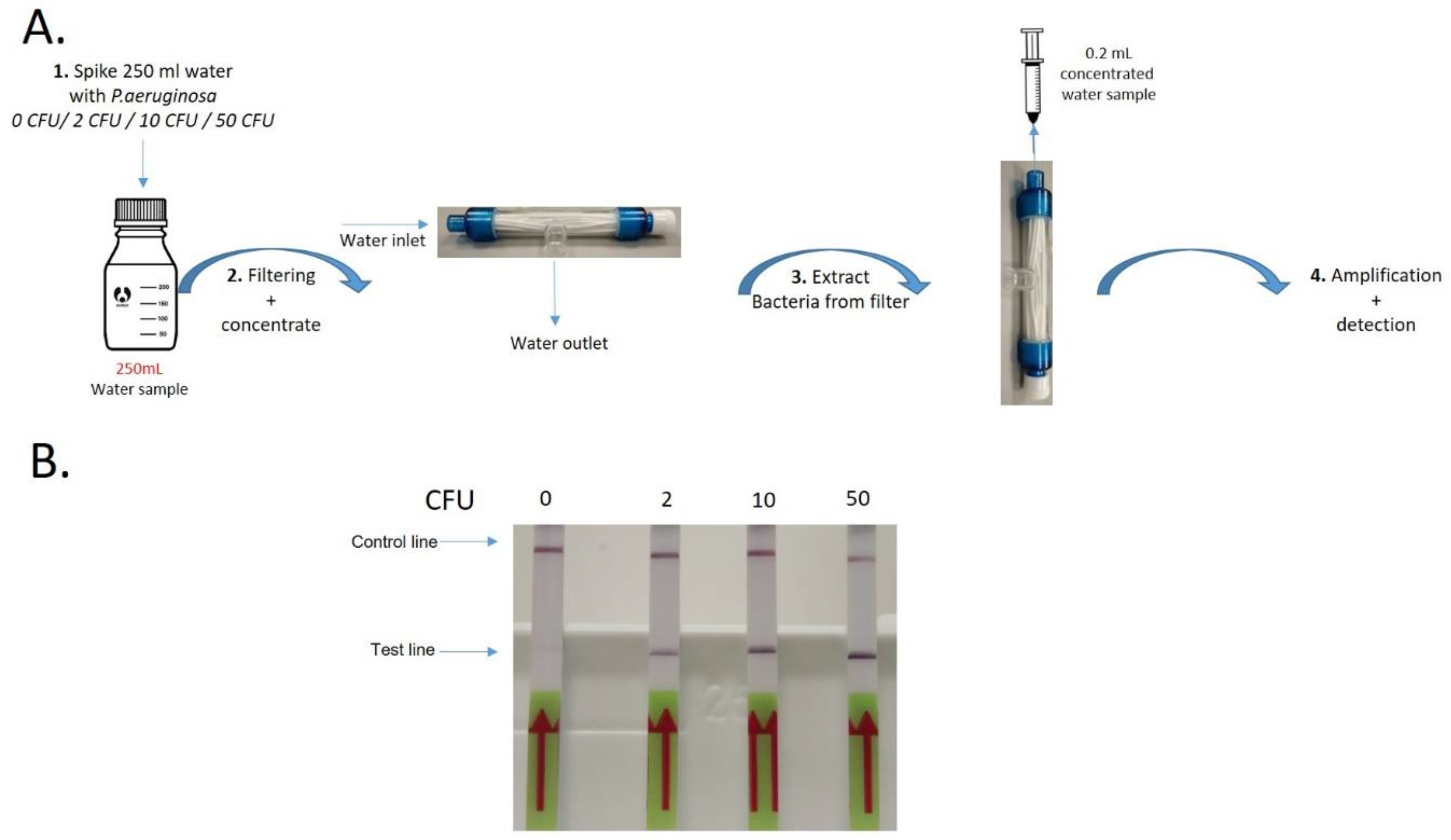
Detection using spiked samples A) Different amount of *P.aeruginosa* are each added to 250 mL of pre-autoclaved ultra-pure water. Standard culture plate count for the *P.aeruginosa* preparation were performed concurrently, so as to ascertain the actual amount of *P.aeruginosa* in the test samples, as per during the test. The spiked water samples are each subjected to a concentration step via a 0.22μm hybrid dead-end/cross flow filter, where the bacteria is captured and subsequently flushed out using a small amount of water/buffer. The concentrated samples are each subjected to the test assay as aforementioned. B) Shown here is the test result and the actual amount of *P.aeruginosa* as per during the day of the test. All the samples that contain *P.aeruginosa* yielded positive test results.

**Table 1:** Overall test performance.

| Reference | Sensitivity (%) | Specificity (%) | PPV (%) | NPV (%) |
| --- | --- | --- | --- | --- |
| qPCR | 96.7 | 92.9 | 96.7 | 92.9 |
| Culture | 92.3 | 83.3 | 92.3 | 83.3 |

Based on the test we carried out, we were able to detect the bacteria of interest, down to a limit of <1 CFU per 100 mL sample. Overall, the test performance achieved was highly favourable, with most of the test measurement/indicators returning high performance of over 90 percent. The specificity and NPV derived using the culture method as reference were lower than desired but was remarkably higher than expected (C.J. et al. 2006). Both the RPA and qPCR methods are molecular detection approaches where the bacteria’s DNA are amplified and detected. The culture method, on the other hand, is a completely different approach, and varying degree of differences in results are expected. Factors, such as culturability, viability, amplification inhibition (due to contaminating inhibitors), etc, are expected to affect the test outcome on RPA and culture differently. This could explain why the test result obtained using qPCR as reference was overall better. To further understand the performance of our test method, future studies, utilising actual field samples, encompassing different matrices, will be required.

As a rapid and deployable bacteria detection system and kit, our method can potentially be used for a wide variety of application, such as infection control for healthcare facility and equipment, food safety, rapid clinical diagnostic test (such as sputum, throat or Group B Streptococcus swab test for prenatal care), etc. Currently, our test method is a qualitative test, where it is only able to indicate the presence or absence of the target bacteria, and is unable to quantitate the amount of bacteria present. For bacteria that are ubiquitously present and do not pose significant risk at low concentration, such a test might not be suitable, and further development will be needed. However, it might still be useful as an early warning/surveillance tool for many of these application, to determine if further test are needed. Currently, there are various semi-quantitative method and tools in the market, such as a lateral kit reader, where the intensity of the test bands can be measured and correlate against known standards. This however will not be suitable for endpoint assay, like in our test method, where the amount of product produced are no longer in linearity with time. Other test strategies such as the most probable number method, which utilises a serial dilution, or amplification method using self-quenching fluorescence probe (similar to the TaqMan probe), where amplification can be monitored and measured and quantitated at real-time, are potential directions towards semi-quantitation and/or full-quantitation.

Besides the issue of quantitation, another striking issue with a DNA-based molecular method, like in our case, will be the inability to differentiate live and dead bacteria. From a public health perspective, generally, only viable pathogens are of major concerns. The method described here, in essence, will amplify DNA from both dead and live cells (after lysis) and/or free-floating DNA from the dead targets, already present in the water sample, considering that DNA are generally very stable, even under harsh environment. Future work, such as the inclusion of a DNase pretreatment and/or the use of a cell-impermeable DNA crosslinkers, such as Ethidium monoazide might be useful (Rudi et al. 2005).

## CONCLUSION

In summary, here we presented and evaluated a fully-deployable pathogen detection method, which potentially could be used in the rapid detection of pathogens in many other application. The test method could also be modified and refined in a number of different ways, to suit the different applications.

## CONTRIBUTORS

J-.Y.C., D.P., B.-H.T., E.K., J.T., H.-M.Y., M.L., L.-T.S., Y.-K.L., K.L.: Conception and design. J-.Y.C., Y.-P.T.: Acquisition and interpretation of data. J-.Y.C., H.-M.Y, K.L.: Drafting or revision of the manuscript.

## ACKNOWLEDGEMENTS

This work was supported by the National Research Foundation (Prime Minister’s Office, Singapore) and the Environment and Water Industry Programme Office (Public Utilities Board, Singapore, research grant, 1301-IRIS-27.

## Footnotes

* Patent pending (Singapore Patent Application No. 10201801064R)

† The CFU unit reported here corresponds to the actual cell number (per test), on the assumption that the cells in our preparation are 100 % viable/culturable.

